# Role of Pex31 in metabolic adaptation of the nucleus vacuole junction NVJ

**DOI:** 10.1101/2025.05.29.656872

**Authors:** Marie Hugenroth, Pascal Höhne, Xue-Tong Zhao, Mike Wälte, Duy Trong Vien Diep, Rebecca Martina Fausten, Maria Bohnert

## Abstract

The nucleus vacuole junction NVJ in yeast is a multifunctional contact site between the nuclear ER membrane and the vacuole with diverse roles in lipid metabolism, transfer and storage. Adaptation of NVJ functions to metabolic cues is mediated by a striking remodeling of the size and the proteome of the contact site, but the extent and the molecular determinants of this plasticity are not fully understood. Using microscopy-based screens, we monitored NVJ remodeling in response to glucose availability. We identified Pex31, Nsg1, Nsg2, Shr5, and Tcb1 as NVJ residents. Glucose starvation typically results in an expansion of the NVJ size and proteome. Pex31 shows an atypical behavior, being specifically enriched at the NVJ at high glucose conditions. Loss of Pex31 uncouples NVJ remodeling from glucose availability, resulting in recruitment of glucose starvation-specific residents and NVJ expansion at glucose replete conditions. Moreover, *PEX31* deletion results in alterations of sterol ester storage and a remodeling of vacuolar membranes that phenocopy glucose starvation responses. We conclude that Pex31 has a role in metabolic adaptation of the NVJ.

**SUMMARY STATEMENT:** Using microscopy-based screens in yeast, we identified Pex31, Nsg1, Nsg2, Shr5 and Tcb1 as residents of the nucleus vacuole junction NVJ. Pex31 has a role in NVJ adaptation to glucose availability.

## INTRODUCTION

Eukaryotic cells are equipped with a set of membrane-bounded organelles that provide distinct, biochemically optimized compartments. Organelle functions are coordinated via contact sites, structures at which different organelles are physically linked by tether machineries to mediate selective material transfer, to enable signal transmission, and to locally organize enzymatic activities (Eisenberg-Bord et al., 2016; Scorrano et al., 2019). The cellular contact site landscape is not static. Instead, metabolic alterations and environmental cues promote re-organizations of the extent, composition and functions of contact sites to enable coordinated cellular adaptations (Bohnert, 2020).

A prominent contact site in *Saccharomyces cerevisiae* (from here on: yeast) is the nucleus vacuole junction (NVJ). This structure links the nuclear ER and the vacuole (yeast lysosome) via an interaction of the nuclear ER protein Nvj1 and the vacuole surface protein Vac8 (Pan et al., 2000). The NVJ is a multifunctional contact site. Originally described as a structure involved in a special form of autophagy termed piecemeal microautophagy of the nucleus (Roberts et al., 2003), the NVJ has emerged as a central platform for lipid handling. Several NVJ resident proteins involved in diverse branches of lipid metabolism have been identified, including Tsc13, an enzyme involved in fatty acid elongation (Kohlwein et al., 2001), the fatty acyl-CoA synthetase Faa1 (Hariri et al., 2018), the 3-hydroxy-3-methylglutaryl-coenzyme A (HMG-CoA) reductases Hmg1 and Hmg2 (Rogers et al., 2021), the phosphatidic acid phosphatase Pah1 (Barbosa et al., 2015), and Cvm1 (Bisinski et al., 2022), a protein involved in sphingolipid metabolism. A further group of NVJ proteins is involved in inter-organelle lipid transfer: Osh1 (Elbaz-Alon et al., 2015; Gatta et al., 2015; Kvam and Goldfarb, 2004; Levine and Munro, 2001), Lam6 (Elbaz-Alon et al., 2015; Gatta et al., 2015; Murley et al., 2015), Nvj2 (Liu et al., 2017; Toulmay and Prinz, 2012), and Vps13 (Lang et al., 2015). Mdm1 is an NVJ component that has a role in the localized formation of lipid droplets, ER-derived storage organelles for neutral lipids (Hariri et al., 2018; Henne et al., 2015). Nvj3, a protein with structural similarities to Mdm1, is also an NVJ resident (Hariri et al., 2018; Henne et al., 2015). Two further NVJ proteins related to organelle biogenesis are Pex29 and Pex30 (Ferreira and Carvalho, 2021). These two proteins are part of a structurally related family of ER membrane proteins that comprises three additional members, Pex28, Pex31 and Pex32. Originally, all family members were described as proteins required for regular abundance and morphology of peroxisomes (Vizeacoumar et al., 2003; Vizeacoumar et al., 2004). Later, roles of Pex30 in biogenesis of pre-peroxisomal vesicles (Joshi et al., 2016) and lipid droplets (Choudhary et al., 2020; Joshi et al., 2018; Wang et al., 2018) from the ER membrane were described. In summary, the NVJ houses a set of proteins that fulfill multiple functions in lipid metabolism and transport, and in organelle biogenesis from the ER.

A prominent feature of the NVJ is its remarkable degree of metabolic adaptability. While the NVJ is typically a small structure at nutrient replete conditions, it strongly expands in response to different types of nutrient deprivation (Hariri et al., 2018; Roberts et al., 2003; Tosal-Castano et al., 2021). During glucose deprivation, this adaptation of the NVJ size depends on the NVJ resident protein Snd3 (Tosal-Castano et al., 2021). Importantly, not only the size of the NVJ adapts to metabolic cues, but also its proteome. While some NVJ proteins, such as the main tether pair Nvj1-Vac8, are permanently present at the NVJ, a large fraction of the known NVJ proteins are conditional NVJ residents that are not enriched at the NVJ at nutrient repletion but accumulate at the contact site when cells run out of glucose. Such a glucose deprivation-induced NVJ recruitment has been described for Snd3 (Tosal-Castano et al., 2021), Hmg1/2 (Rogers et al., 2021), Pah1 (Barbosa et al., 2015), Nvj2 (Toulmay and Prinz, 2012), Faa1 (Hariri et al., 2018), Vps13 (Lang et al., 2015), and Pex29/30 (Ferreira and Carvalho, 2021). However, remodeling of the NVJ proteome has not been systematically studied, and the molecular determinants of the proteomic adaptation of the NVJ have been largely unclear.

Here, we used systematic microscopy-based approaches to compare the NVJ at glucose-replete and -restricted conditions and identified five additional NVJ proteins, the permanent NVJ resident Shr5 and the conditional residents Nsg1, Nsg2, Tcb1, and Pex31. While Nsg1, Nsg2, and Tcb1 belong to the large group of conditional NVJ residents that accumulate at low-glucose conditions, Pex31 is unique as it is enriched at the NVJ at high glucose conditions, but not during glucose starvation. *PEX31* deletion results in a dysregulation of NVJ remodeling: Low glucose-specific NVJ features, such as NVJ enlargement and expansion of the NVJ proteome, occur independent of glucose availability in Δ*pex31* cells. We conclude that Pex31 has a role in NVJ adaptation to metabolic cues.

## RESULTS

### A microscopy-based screen identifies residents of the nucleus vacuole junction NVJ

Alterations in glucose availability result in pronounced adaptations of the NVJ proteome (Barbosa et al., 2015; Ferreira and Carvalho, 2021; Hariri et al., 2018; Lang et al., 2015; Rogers et al., 2021; Tosal-Castano et al., 2021; Toulmay and Prinz, 2012), but the extent of these remodeling processes and the underlying molecular determinants are only partially understood. To gain a broad view on NVJ remodeling, we performed a microscopy-based screen. We assembled a mutant collection of 337 strains expressing N-terminally GFP tagged proteins from a genome-wide mutant library (Weill et al., 2018; Yofe et al., 2016). The collection comprised known NVJ residents and their paralogs as well as proteins involved in lipid metabolism and lipid transport. We then used an automated mating approach (Cohen and Schuldiner, 2011; Tong and Boone, 2006) to introduce the NVJ marker Nvj1-Cherry into this mutant collection. In a parallel approach, we used the vacuolar membrane marker Zrc1-Cherry as an alternative way to visualize the vacuole-nuclear ER interface without manipulation of an integral NVJ protein. We complemented these mutant collections with select manually created strains expressing C-terminally mNeonGreen (mNG) tagged proteins (Fig. 1A). We also included a strain expressing a Vps13 variant internally tagged with GFP that was previously described as a functional NVJ protein (Lang et al., 2015). All strains were analyzed by automated microscopy at two different metabolic conditions: Cells were either grown overnight on glucose replete medium, back-diluted to fresh glucose replete medium and further grown to logarithmic growth phase (Fig. 1A, High D), or precultured on glucose replete medium overnight, and then incubated for four hours in glucose restriction medium containing only 0.001% glucose (Fig. 1A, D restr.). GFP/mNG-tagged proteins were then classified according to their localization at the NVJ (Fig. 1B, S1A-D). Overall, we identified 20 NVJ residents, five of which have, to our knowledge, not been described before. Eight proteins were present at the NVJ at both high and low glucose conditions (Fig. 1B, permanent NVJ residents), 11 accumulated at the NVJ specifically at glucose restriction (Fig. 1B, conditional NVJ resident at glucose restriction), and one was found enriched at the NVJ selectively at the glucose replete condition (Fig. 1B, conditional NVJ residents at high glucose). The five previously unknown NVJ proteins (Fig. 1B, grey boxes) were re-analyzed at the high glucose condition and at the acute glucose restriction condition used in the screen. Additionally, we included a protocol for a gradual form of glucose starvation, by growing cells in medium supplemented with 2% glucose for 20-22 hours into early stationary phase, resulting in glucose exhaustion from the medium (D exh.). Shr5 has previously been described as an ER-resident protein involved in palmitoylation and correct targeting of Ras2 (Dong et al., 2003; Jung et al., 1995; Lobo et al., 2002). We detected this protein at the NVJ at all three conditions analyzed (Fig. 1C, S1C), classifying it as a permanent NVJ resident (Fig. 1B), similar to the known NVJ proteins Nvj1, Nvj3, Mdm1, Osh1, Lam6, Tsc13, and Vac8 (Fig. S1A). Nsg1 and Nsg2 are two homologs of the mammalian INSIG (insulin-induced gene) proteins that are involved in regulation of sterol levels (Flury et al., 2005). Nsg1 and 2 have been found to bind to the sterol-sensing domain of the HMG-CoA reductase Hmg2, thereby stabilizing this enzyme that mediates the rate-limiting enzymatic step in the mevalonate pathway (Basson et al., 1986; Flury et al., 2005). Both Hmg2 and its homolog Hmg1 have in the past been shown to localize to the NVJ specifically during glucose starvation (Rogers et al., 2021). We find that Nsg1 and Nsg2 show a similar behavior by accumulating at the NVJ specifically upon glucose exhaustion or restriction (Fig. 1D, S1D). A further protein that we found enriched at the NVJ in response to the low glucose conditions, but not at high glucose, is Tcb1 (Fig. 1D, S1D). Tcb1 is part of the family of tricalbin proteins, which have been identified as residents of the contact site between the ER and the plasma membrane (Collado et al., 2019; Creutz et al., 2004; Manford et al., 2012; Toulmay and Prinz, 2012). In summary, we identify Tcb1, Nsg1 and Nsg2 as NVJ proteins preferentially enriched at the contact at low glucose (Fig. 1B), similar to the previously described NVJ residents Hmg1, Hmg2, Snd3, Nvj2, Cvm1, Vps13, Pex29 and Pex30 (Fig. S1B). The latter two proteins are part of a structurally related protein family (Vizeacoumar et al., 2003; Vizeacoumar et al., 2004). Interestingly, we identified a further member of this family, Pex31, at the NVJ, however, this protein showed an inverse behaviour, being enriched at the NVJ at glucose repletion, but not at glucose restriction or exhaustion (Fig. 1E). We analyzed the behavior of this high glucose-specific NVJ resident in more detail and detected a colocalization of Pex31 foci with Nvj1-Cherry in over 40% of cells at glucose repletion (Fig. 1F). We created a strain expressing Pex31-HA from its genomic locus under control of its own promoter and analyzed Pex31 protein levels. We found that Pex31 abundance dropped steeply at glucose restriction as well as exhaustion conditions (Fig. S1E).

**Figure 1.**
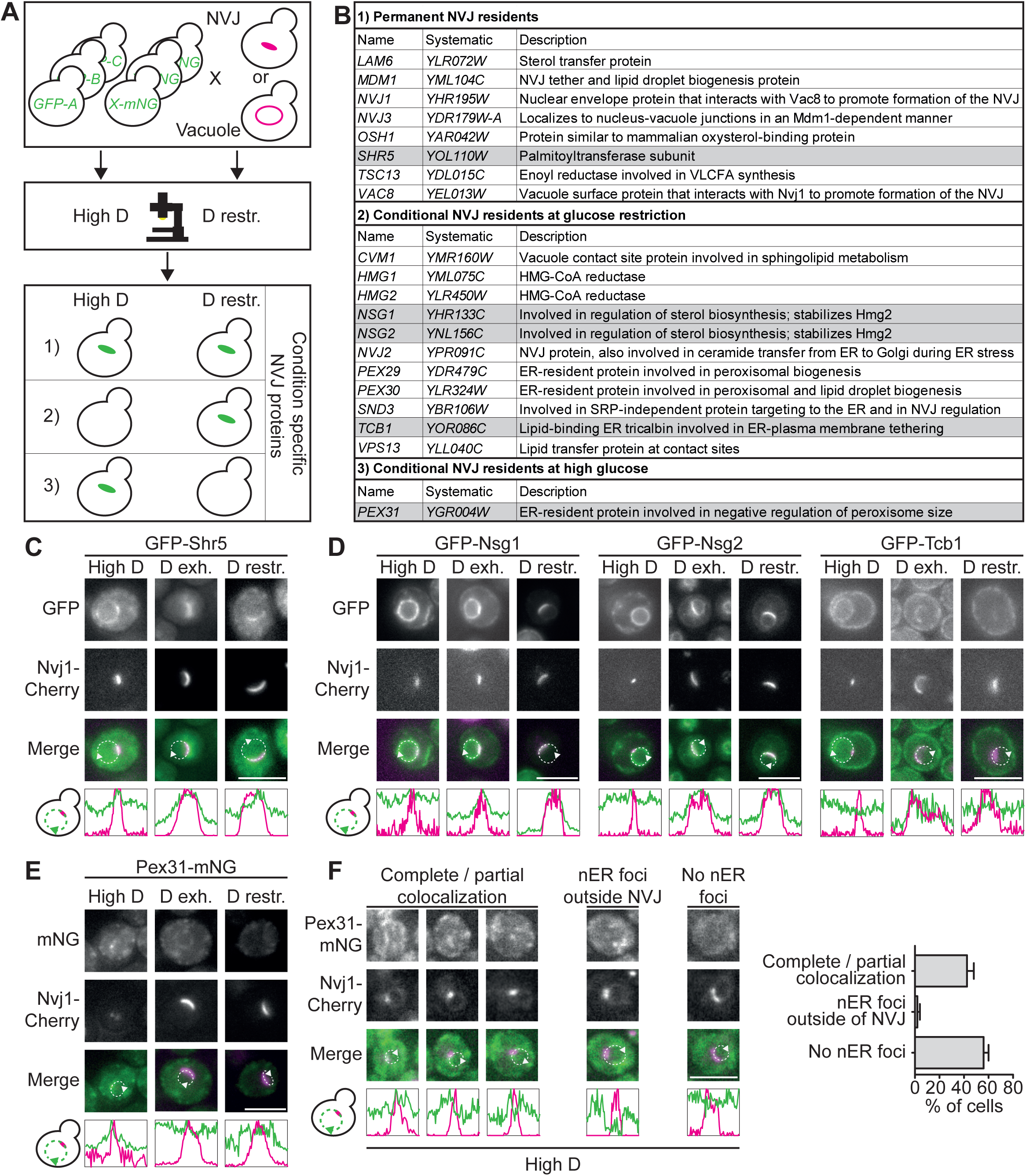
A microscopy-based screen identifies Pex31, Nsg1, Nsg2, Shr5, and Tcb1 as residents of the nucleus vacuole junction. (A) Schematic depiction of a screen searching for NVJ residents at glucose replete and restricted conditions. Nvj1-Cherry or Zrc1-Cherry were introduced as markers of the NVJ into a collection of strains expressing N-terminally GFP- or C-terminally mNeongreen (mNG)-tagged proteins related to lipid handling. Strains were imaged at glucose replete (High D) and glucose restricted (D restr.) conditions and categorized according to their localization to the NVJ: (1) Permanent NVJ residents displaying NVJ localization at both conditions, (2) conditional NVJ residents localizing to the NVJ upon glucose restriction, and (3) conditional NVJ residents localizing to the NVJ at glucose replete conditions. (B) List of hits from screen in (A). Previously described NVJ residents in white, additional NVJ residents in grey. (C) A strain expressing GFP-Shr5 and Nvj1-Cherry was grown at indicated conditions. Arrowhead in line graphs depicts end of the line graph. Brightness and contrast were adjusted individually. Scale bar, 5 µm. (D) Cells co-expressing GFP-Nsg1, GFP-Nsg2 and GFP-Tcb1 with Nvj1-Cherry were grown at indicated conditions. Brightness and contrast were adjusted individually. Scale bar, 5 µm. (E) Pex31-mNG Nvj1-Cherry cells were grown at glucose replete (High D), glucose exhausted (D exh.) and glucose restricted (D restr.) conditions. Brightness and contrast were adjusted individually. Scale bar, 5 µm. (F) Cells from (E) at glucose replete conditions were categorized based on Pex31-mNG localization in reference to the Nvj1-Cherry signal. Example images of each category are shown on the left. Scale bar, 5 µm. Percentage of each category is depicted on the right as mean ± SEM. N=50 cells; n=3.

In summary, we have identified Shr5, Nsg1, Nsg2, Tcb1 and Pex31 as NVJ residents, of which Pex31 is uniquely enriched at the NVJ at the high glucose condition.

### Pex31 limits the NVJ proteome at high glucose

The identification of Pex31 as an atypical NVJ resident with a preference for high glucose conditions prompted us to follow up on this protein. To assess the impact of Pex31 on the NVJ, we introduced a Δ*pex31* allele into a mutant collection expressing N-GFP protein variants (Weill et al., 2018; Yofe et al., 2016) by automated mating (Cohen and Schuldiner, 2011; Tong and Boone, 2006). As in the initial screen (Fig. 1A), the collection was complemented with manually created strains expressing selected C-terminally mNG-tagged proteins. We then cultured both the new mutant collection carrying the Δ*pex31* allele and the unaltered parent library on high glucose medium to logarithmic growth phase, the condition where typically only the permanent NVJ proteins accumulate at the contact site. We analyzed both mutant collections by automated microscopy and compared localization of the tagged proteins (Fig. 2A). We found that localization of the permanent NVJ residents was unaffected by *PEX31* deletion (Fig. 2B, top). In contrast, we observed an effect of the *PEX31* deletion on part of the conditional NVJ proteome. Pex29, Pex30, Nsg1, Hmg2 and Nvj2, which typically accumulate at the NVJ selectively at low glucose conditions (Fig. 1B-D), showed an enrichment at the NVJ on high glucose medium in response to the *PEX31* deletion (Fig. 2B, labeled in gray). We re-created all strains manually to re-assess and quantify localization of the five proteins. Proteins of interest were marked with a green fluorescent marker and either the vacuole was visualized by CMAC staining (labeling the vacuole lumen) or the NVJ was marked by genomic expression of Nvj1-Cherry (Fig. 2C-G, left). All five proteins were found enriched at the NVJ in Δ*pex31* cells (Fig. 2C-G, right), confirming the results of the screen.

**Figure 2.**
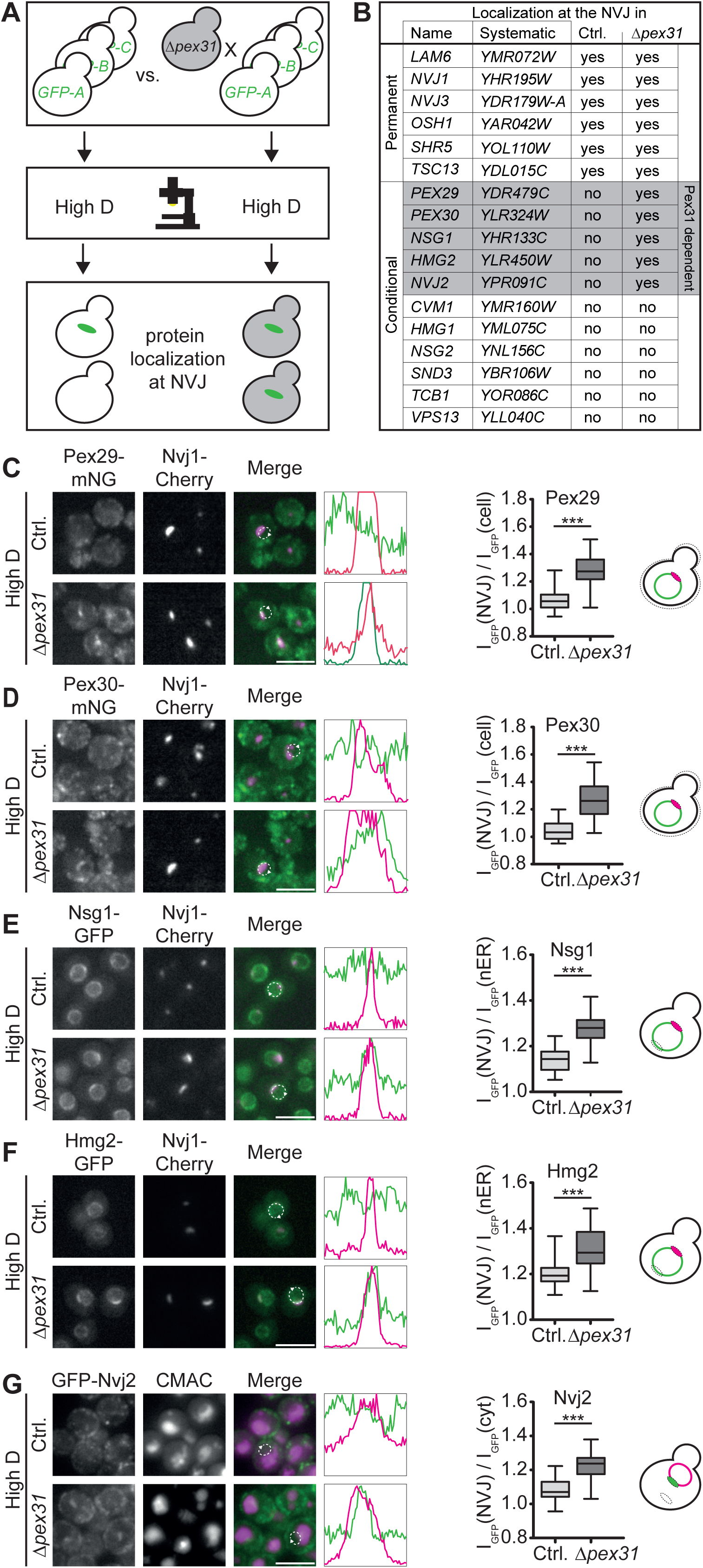
**Loss of Pex31 alters NVJ composition at glucose replete conditions**. (A) Schematic representation of a screen comparing protein localization in the presence and absence of *PEX31*. Automated mating was used to delete *PEX31* in a genome-wide collection of strains expressing N-terminally GFP-tagged proteins under control of a *NOP1* promotor. The resulting mutant collection was imaged alongside the parent collection at glucose replete conditions and all strains were assessed for GFP signal at the NVJ. (B) List of hits from the screen. Marked in grey, proteins re-localizing to the NVJ upon deletion of *PEX31* at glucose rich conditions. (C) Left: Pex29-mNG Nvj1-Cherry (Ctrl.) and Pex29-mNG Δ*pex31* Nvj1-Cherry cells grown on glucose replete conditions. Right: Enrichment of GFP signal at the NVJ (I_GFP_(NVJ)) over whole cell GFP signal (I_GFP_(cell)) was quantified. Data in box plots depicting the median and the whiskers depicting the minimum to maximum values. Compared by t-test. ***p < 0.001. N=30 cells, n=3. (D) Left: Indicated cells grown in glucose replete conditions. Scale bar, 5 µm. Right: Quantification as in (C). (E) Left: Indicated strains grown in glucose replete conditions. Right: Enrichment of GFP signal at the NVJ (I_GFP_(NVJ)) over nuclear ER signal at the opposite site (I_GFP_(nER)) was quantified. Data depicted in box plots showing the median and the whiskers depicting the minimum to maximum values. Compared by t-test. ***p < 0.001. N=30 cells, n=3. (F) Left: Hmg2-GFP and Nvj1-Cherry co-expressed in control (Ctrl.) and Δ*pex31* cells grown in glucose replete conditions. Right: Quantification as in (E) (G) Left: GFP-Nvj2 (Ctrl.) and GFP-Nvj2 Δ*pex31* cells were stained with the vacuole lumen dye CMAC and imaged in glucose replete conditions. Scale bar, 5 µm. Right: Enrichment of GFP signal at the NVJ (I_GFP_(NVJ)) over cytosolic GFP signal (I_GFP_(cyt)) was quantified. Data represented in box plots showing the median and the whiskers depicting the minimum to maximum values. Compared by t-test. ***p < 0.001. N=30 cells, n=3.

In summary, Pex31 is a resident of high glucose NVJs, which typically have a limited proteome, and *PEX31* deletion results in an expansion of the high glucose NVJ proteome in a manner that partially phenocopies low glucose conditions.

### Loss of *PEX31* promotes NVJ enlargement independent of glucose availability

Glucose starvation not only causes an expansion of the NVJ proteome (Fig. 1B), but also an increase of the overall NVJ size (Hariri et al., 2018; Roberts et al., 2003; Tosal-Castano et al., 2021). Knowing that *PEX31* deletion partially phenocopies expansion of the NVJ proteome independent of glucose availability (Fig. 2), we asked if it also affects the size of the NVJ in a similar manner. As an estimate of NVJ size, we quantified the area of Nvj1-Cherry foci at different conditions (Fig. 3A, B). Consistent with previous reports, we observed an expansion of the NVJ in response to low glucose conditions in control cells (1.55 µm² mean NVJ area at glucose exhaustion versus 1.06 µm² at high glucose) (Fig. 3A, B). Furthermore, we found that *PEX31* deletion led to a pronounced NVJ enlargement at high glucose (1.58 µm²). Interestingly, the average size of Δ*pex31* NVJs at high glucose was comparable to the size of control NVJs at glucose exhaustion (Fig. 3A, B). No further expansion of Δ*pex31* NVJs was observed upon glucose exhaustion (Fig. 3A, B), suggesting that NVJs in Δ*pex31* cells have glucose starvation-like features independent of glucose availability.

**Figure 3.**
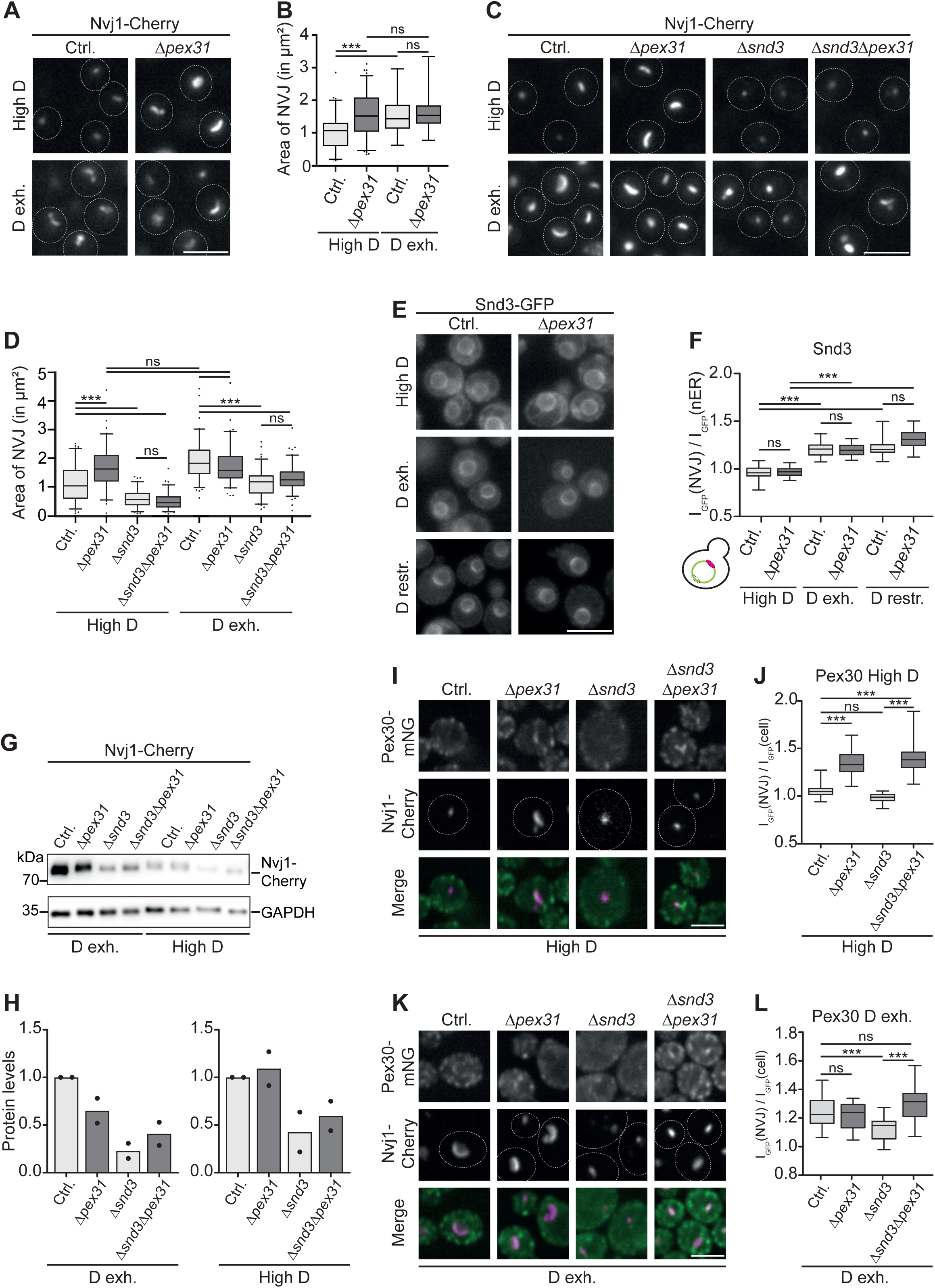
**Deletion of *PEX31* promotes NVJ expansion at high glucose conditions**. (A) Control and Δ*pex31* cells expressing Nvj1-Cherry were grown on either High D or D exh. conditions. Scale bar, 5 µm. (B) Quantification of Nvj1-Cherry signal from (A). Area of NVJ in µm² is depicted as boxplots (5-95 percentile). Compared by one-way ANOVA. ns, not significant; ***p < 0.001. N=50 cells, n=3. (C) Indicated strains expressing Nvj1-Cherry in High D or D exh. conditions. Scale bar, 5 µm. (D) Quantification of Nvj1-Cherry signal from (C). Area of NVJ in µm² is depicted as boxplots (5-95 percentile). Compared by one-way ANOVA. ns, not significant; ***p < 0.001. N=50 cells, n=3. (E) Snd3-GFP control and Snd3-GFP Δ*pex31* cells grown in indicated conditions. Scale bar, 5 µm. (F) Ratio of Snd3-GFP signal at NVJ (I_GFP_(NVJ)) to nuclear ER (I_GFP_(nER)) at the opposite site. Data in graphs represented in box plots showing the median and the whiskers depicting minimum to maximum value. Compared by one-way ANOVA. ns, not significant. ***p < 0.001 (G) Control (Ctrl.), Δ*pex31,* Δ*snd3*, Δ*snd3*Δ*pex31* cells expressing Nvj1-Cherry were grown under indicated conditions and subjected to SDS-PAGE and subsequent western blotting. (H) Quantification of Western Blot shown in (G). Nvj1-Cherry intensity was normalized to GAPDH intensity and compared to respective control. n=2. (I) Cells expressing Pex30-mNG and Nvj1-Cherry in either control (Ctrl.), Δ*pex31*, Δ*snd3* or Δ*snd3*Δ*pex31* background. Grown in high glucose. Scale bar, 5 µm. (J) Ratio of Pex30-mNG signal under glucose replete conditions at NVJ (I_GFP_(NVJ)) to cellular GFP signal (I_GFP_(cell)) depicted as box plots (minimum to maximum value). Compared by one-way ANOVA. ns, not significant; ***p < 0.001. N=50 cells, n=3. (K) Analysis as in (I) with the difference that cells were grown in glucose exhaustion (D exh.). (L) Ratio of Pex30-mNG signal under glucose exhaustion at NVJ (I_GFP_(NVJ)) to cellular GFP signal (I_GFP_(cell)) depicted as box plots (minimum to maximum value). Compared by one-way ANOVA. ns, not significant; ***p < 0.001. N=50 cells, n=3.

The conditional NVJ resident Snd3 has been identified as an important player in controlling NVJ size. *SND3* deletion causes an overall reduction in NVJ size across metabolic conditions and a block of NVJ expansion at glucose exhaustion (Tosal-Castano et al., 2021). Consistent with these findings, we observed that *SND3* deletion resulted in a decrease in the NVJ size at high glucose (0.61 µm² average NVJ area in *Δsnd3* compared to 1.12 µm² in control cells) (Fig. 3C, D). When we simultaneously deleted *SND3* and *PEX31*, we found that *Δpex31Δsnd3* double mutant cells had small NVJs (0.55 µm²), comparable to those observed in *Δsnd3* single deletion cells (Fig. 3C, D), showing that Snd3 is required for NVJ expansion in response to loss of Pex31.

Based on the notion that loss of Pex31 promotes accumulation of conditional NVJ proteins, we next asked if the NVJ expansion observed in *Δpex31* cells was mediated by recruitment of the NVJ regulator Snd3 to the NVJ. However, while Snd3 accumulated at the NVJ in response to low glucose conditions as reported (Tosal-Castano et al., 2021), Snd3 distribution was not affected by loss of Pex31 (Fig. 3E, F), consistent with the results of our screen for *PEX31* dependent NVJ residents (Fig. 2B).

Loss of Snd3 has been reported to affect the NVJ by triggering degradation of the main NVJ tether Nvj1 (Tosal-Castano et al., 2021). We therefore assessed Nvj1-Cherry protein levels in Δ*pex31*, *Δsnd3* and *Δpex31Δsnd3* cells by western blot at high glucose and glucose exhaustion conditions. As expected, Nvj1 levels were consistently higher at glucose exhaustion (Fig. 3G).

We found that at high glucose conditions, Nvj1 levels were unaffected by *PEX31* deletion (Fig. 3G, H), showing that NVJ expansion in Δ*pex31* cells at this condition was not mediated by alterations in the amount of this tether protein. Deletion of *SND3* on the other hand resulted in reduced Nvj1 levels at both conditions (Fig. 3G, H). This reduction in Nvj1 levels did not only occur in the *Δsnd3* single mutant, but also in *Δpex31Δsnd3* double mutant cells (Fig. 3G, H). In summary, *PEX31* deletion does not promote Snd3 accumulation at the NVJ, suggesting that NVJ expansion in *Δpex31* cells is not directly driven by Snd3 recruitment. However, deletion of *SND3* leads to a partial loss of the key NVJ tether Nvj1, resulting in a general block of NVJ expansion not only upon glucose exhaustion (Tosal-Castano et al., 2021), but also in response to *PEX31* deletion.

Finally, we asked if Snd3 is required for remodeling of the NVJ proteome upon *PEX31* deletion. To test this, we assessed localization of Pex30 in Δ*pex31*, *Δsnd3* and *Δpex31Δsnd3* double mutant cells. We found that Pex30 accumulated at the NVJ in Δ*pex31* cells at high glucose both in the presence and in the absence of Snd3 (Fig. 3I, J). This shows not only that Snd3 is dispensable for Pex30 re-localization in Δ*pex31* cells but also suggests that Pex31-dependent modulation of the NVJ proteome does not directly depend on the increase in contact site size, as Pex30 was efficiently recruited to the small NVJs of *Δpex31Δsnd3* double mutant cells. We also analyzed Pex30 distribution at glucose exhaustion and found that it accumulated at the NVJ as reported (Ferreira and Carvalho, 2021) (Fig. 3K, L). Of note, Pex30 was lost from NVJs in Δ*snd3* cells, and this effect was reversed by additional deletion of *PEX31*, further supporting the notion that loss of Pex31 promotes an expansion of the NVJ proteome (Fig. 3K, L).

In summary, loss of Pex31 leads to an increased NVJ size at high glucose conditions, phenocopying the effects of glucose starvation. Both expansion of the NVJ proteome and NVJ enlargement via *PEX31* deletion are not directly mediated via the NVJ regulator Snd3. However, loss of the Nvj1 tether protein upon *SND3* deletion blunts NVJ expansion.

### Δ*pex31* cells show multiple hallmarks of starvation

Loss of Pex31 results in alterations of both NVJ proteome and size that are reminiscent of NVJ features typically observed at glucose starvation. This prompted us to test if Δ*pex31* cells display additional hallmarks of glucose starvation related to the vacuole.

A characteristic feature of glucose exhausted or restricted cells is an alteration in vacuolar morphology. While cells often have multilobed vacuoles at high glucose conditions, low glucose typically promotes formation of a single large, spherical vacuole (Armstrong, 2010; Li and Kane, 2009). To assess vacuolar morphology, we stained control and Δ*pex31* cells at glucose replete, restricted and exhausted conditions with the vacuolar lumen dye CMAC. As expected, control cells showed a mixed phenotype with both multilobed and single vacuoles at glucose repletion, while vacuoles typically fused to one large structure at low glucose conditions (Fig. 4A). In contrast, Δ*pex31* cells often showed large single vacuoles independent of glucose availability, phenotypically mimicking low-glucose conditions in the presence of high glucose (Fig. 4A). To quantify this effect, the vacuolar membrane marker Vph1-mKate2 was expressed in control and Δ*pex31* cells to optimally resolve single vacuoles. Quantification of cells with one vacuole versus cells with two or more vacuoles confirmed a pronounced shift toward single large vacuoles at high glucose in Δ*pex31* cells (Fig. 4B).

**Figure 4.**
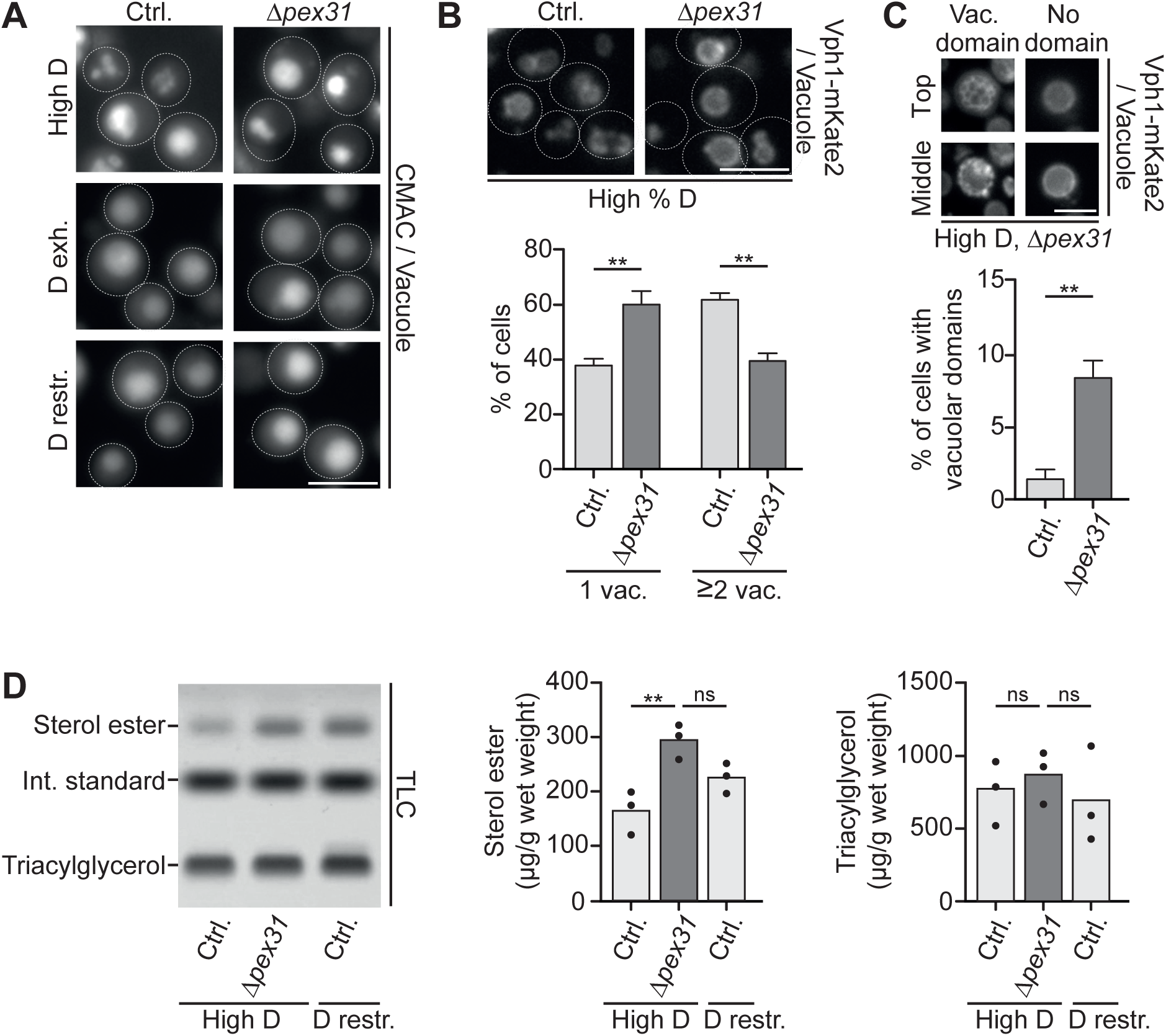
**Absence of Pex31 induces glucose starvation-like phenotypes of lipid storage and vacuolar organization**. (A) Control and Δ*pex31* cells stained with CMAC for visualization of vacuoles. Grown at indicated conditions. Scale bar, 5 µm. (B) Top: Control and Δ*pex31* cells expressing Vph1-mKate2 as a vacuolar marker in glucose replete conditions. Scale bar, 5 µm. Bottom: Mean of percentage of cells showing one vacuole (1 vac.) or at least two vacuoles (≥2 vac.). Compared by t-test. **p < 0.01. N=50 cells, n=3. (C) Top: Example images of cells with vacuolar domains (Vac. domain) and without (No domain) in Δ*pex31* cells. Z-stacks of Vph1-mKate2 expressing cells were analyzed. Bottom: Percentage of cells with vacuolar domains in glucose replete conditions. Data represented as mean ± SEM. Compared by t-test. **p < 0.01. N=50 cells, n=3. (D) Left: High performance thin layer chromatography (HPTLC) plate of lipid extracts from control (Ctrl.) and Δ*pex31* cells in high glucose (High D) or glucose restriction (D restr.) showing neutral lipid separation. Int. standard = Internal standard (cholesteryl formate). Right: Quantification of sterol ester and triacylglycerol in µg/g wet weight. ns, not significant; **p < 0.01. n=3.

Glucose exhaustion has also been found to promote a lateral re-organization of the vacuolar membrane. Liquid ordered, likely sterol-rich, and liquid disordered membrane domains are formed, which can be visualized via Vph1-mKate2 that specifically segregates into the liquid disordered membrane regions (Toulmay and Prinz, 2013; Wang et al., 2014). Consistent with previous observations, we found that these domains were very rare at glucose repletion in control cells (around 1%). In contrast, around 8% of Δ*pex31* cells showed vacuolar membrane domains at glucose repletion (Fig. 4C). In conclusion, Δ*pex31* vacuoles at high glucose conditions show multiple features of glucose starvation.

Finally, we asked if Δ*pex31*-dependent alterations had consequences on NVJ function. It has been found that accumulation of the HMG-CoA reductases Hmg1 and Hmg2 at the NVJ during glucose restriction promotes flux through the mevalonate pathway and sterol ester synthesis (Rogers et al., 2021). We find that *PEX31* deletion drives not only glucose-independent accumulation of Hmg2 at the NVJ, but also a similar re-localization of the Hmg2 stabilizer Nsg1 (Fig. 2B, 2E, 2F). To test if this Pex31-dependent re-organization reflects on the cell’s lipidome, we used high-performance thin-layer chromatography to analyze the neutral lipid content of control and Δ*pex31* cells at glucose replete conditions. We found that *PEX31* deletion indeed resulted in an increase in sterol ester levels at glucose repletion, while levels of triglycerides were unaffected (Fig. 4D). Similar to our findings on the size (Fig. 3) and composition (Fig. 2) of the NVJ, sterol ester levels Δ*pex31* cells at glucose repletion were comparable to those observed in glucose restricted control cells (Fig. 4D).

In summary, loss of Pex31 not only provokes a re-organization of the NVJ reminiscent of alterations that typically occur at low-glucose conditions, but also further glucose starvation-related adaptations of vacuole remodeling and neutral lipid storage.

## DISCUSSION

Here, we have used microscopy-based high-content screens to characterize the NVJ at glucose replete and starved conditions and have identified permanent and condition-specific contact site residents. We find that one of these residents, Pex31, has a role in metabolic adaptation of the size and composition of the NVJ. The contact site generally has a limited size and proteome at high glucose conditions. Upon glucose starvation, the NVJ size increases and the contact site becomes populated by additional residents. Intriguingly, Pex31 shows the inverse behavior, being enriched at the small NVJ structures observed at glucose repletion and being lost from the contact at starvation. Furthermore, *PEX31* deletion uncouples NVJ expansion from metabolic cues: In Δ*pex31* cells, the NVJ expands at high glucose, comparable to the size typically observed during glucose starvation. At the same time, a set of normally glucose starvation-specific contact site residents populate the Δ*pex31* NVJ irrespective of high glucose availability. Overall, these findings are consistent with a model where Pex31 has a role in restricting NVJ size and composition at glucose repletion.

We assessed the relationship of Pex31 to Snd3, a further NVJ modulator. Deletion of *SND3* has been shown to result in destabilization of the key NVJ tether Nvj1 and a block in NVJ expansion in response to glucose starvation (Tosal-Castano et al., 2021). We find that loss of Snd3 and the concomitant decrease in Nvj1 tether levels broadly blunts NVJ size expansion, not only at glucose starvation, but also in Δ*pex31* cells. However, expansion of the NVJ proteome in Δ*pex31* cells was unaffected by *SND3* deletion, showing that NVJ remodeling by Pex31 is not directly mediated by Snd3.

Pex31 is part of a structurally related protein family, which, besides Pex31 itself, comprises Pex28, Pex29, Pex30 and Pex32. While the exact molecular roles of the Pex28/29/30/31/32 proteins are unclear, it is emerging that their functions are related to the maintenance of ER membrane homeostasis. All family members are ER-integral proteins anchored in the membrane by an N-terminal reticulon homology domain (RHD) (Joshi et al., 2016), which is similar to the RHDs of the membrane shaping reticulon proteins (Voeltz et al., 2006). Pex30 and Pex31 overexpression indeed has been found to restore viability of a mutant simultaneously lacking the reticulons Rtn1 and Rtn2, the reticulon-binding protein Yop1, and the lipid metabolism regulator Spo7, an effect that has been suggested to be linked to the ER membrane shaping ability of the Pex30/31 RHD (Joshi et al., 2016). C-terminal to the RHD, all members of the family comprise an additional Dysferlin domain. These domains bind to phosphatidic acid in vitro, and loss of the Pex28/29/30/31/32 proteins results in aberrant phosphatidic acid distribution within the ER membrane (Ferreira et al., 2025; House et al., 2025). All Pex28/29/30/31/32 proteins affect formation of ER-derived organelles. Individual loss of each family member results in altered abundance and morphology of peroxisomes (Vizeacoumar et al., 2003; Vizeacoumar et al., 2004). Pex30 has been found to have a role in the organization of specialized ER membrane subdomains and in the formation of pre-peroxisomal vesicles from these domains (Joshi et al., 2016). Additionally, Pex30 is involved in formation of lipid droplets from the ER membrane (Choudhary et al., 2020; House et al., 2025; Joshi et al., 2018; Wang et al., 2018). Immunoprecipitation revealed that Pex30 forms different heterooligomeric complexes with other family members that localize to distinct organelle contact sites: Pex28/30/32 complexes that reside at ER-peroxisome contacts, and Pex29/30 complexes, which localize to the NVJ at glucose starvation (Ferreira and Carvalho, 2021). Intriguingly, the same study revealed that Pex31 is uniquely excluded from complex formation with Pex30 (Ferreira and Carvalho, 2021), raising the question about the functional relation of this protein. We find that Pex31 is also an NVJ resident, however, its response to metabolic states and its effect on the contact site are inverse. While Pex29/30 localizes to the NVJ specifically at glucose starvation, Pex31 is enriched at this contact site at glucose repletion. Furthermore, while Pex29/30 are required for NVJ expansion (Ferreira and Carvalho, 2021; Ferreira et al., 2025), we find that Pex31 counteracts the expansion of the NVJ size and proteome, suggesting an antagonistic effect of Pex29/30 and Pex31. Interestingly, *PEX31* deletion not only phenocopies a state of starvation in terms of NVJ organization. Instead, further hallmarks of starvation are induced, including a shift from multilobed to single, round vacuoles, a re-organization of the vacuole membrane into liquid ordered and disordered domains, and an increase in the storage of sterol esters. The exact mechanistic basis of this starvation-like organelle reprogramming via Pex31 is an important topic for future studies.

More broadly, we have expanded our view on the proteome of the NVJ, offering future directions for the analysis of this central organelle interface. The finding that the Hmg2 stabilizing proteins Nsg1 and Nsg2 accumulate at the NVJ at glucose starvation opens interesting directions for the regulation of sterol metabolism. Nsg1 and Nsg2 are the yeast homologs of the mammalian insulin-induced gene (INSIG) proteins (Sever et al., 2003; Yang et al., 2002). Nsg1 and Nsg2 stabilize Hmg2 by preventing its degradation via ER associated degradation (Flury et al., 2005). We find that not only glucose starvation but also manipulation of the NVJ by loss of Pex31 induce a re-localization of both Hmg2 and Nsg1 to the NVJ. The details of how this mechanistically relates to the increased sterol ester accumulation that we detect in Δ*pex31* cells remain to be determined. We also detect Shr5 at the NVJ, a subunit of a palmitoyl transferase that together with Erf2 palmitoylates Ras nutrient signaling GTPases, regulating their membrane localization and activation (Dong et al., 2003; Jung et al., 1995; Lobo et al., 2002; Zhao et al., 2002). Investigating the role of Shr5 at the NVJ and potential implications for Ras signaling is an interesting topic for the future. Finally, we detect Tcb1 at the NVJ upon glucose starvation, a member of the Tcb1/2/3 tricalbin family and a well-characterized ER-plasma membrane contact site resident that acts as tether and lipid transfer protein (Collado et al., 2019; Creutz et al., 2004; Hoffmann et al., 2019; Manford et al., 2012; Toulmay and Prinz, 2012). Interestingly, *NVJ1* shows negative genetic interactions with the tricalbins (Hoffmann et al., 2019). This suggests partially redundant functions of ER-plasma membrane contacts and the NVJ. Alternatively, the finding that Tcb1 can re-localize to the NVJ opens the possibility for a more direct mechanism underlying this genetic link. The exact role of Tcb1 at the NVJ remains elusive, but it is tempting to speculate that the dual localization to ER contact sites with the plasma membrane and the vacuole might serve a regulatory role. Similar dual localizations to distinct contact sites have been described e.g. for Vps13 (Lang et al., 2015), Nvj2 (Liu et al., 2017; Toulmay and Prinz, 2012), Osh1 (Elbaz-Alon et al., 2015; Gatta et al., 2015; Kvam and Goldfarb, 2004; Levine and Munro, 2001), and members of the Lam protein family (Elbaz-Alon et al., 2015; Gatta et al., 2015; Murley et al., 2015).

In conclusion, we have in this work identified multiple additional NVJ residents, and a regulator of proteomic NVJ adaptation to metabolic cues, opening new directions toward a mechanistic understanding of the functional plasticity of this intriguing organelle interface.

## MATERIALS AND METHODS

### Yeast strains and growth conditions

*Saccharomyces cerevisiae* strains used in this study were derived from a synthetic genetic array (SGA) compatible strain (Breslow et al., 2008) and are described in Table S1. Cells were transformed with PCR products using a method including lithium-acetate, polyethylene glycol, and single stranded DNA (Gietz and Woods, 2006; Longtine et al., 1998). Primers for genetic modifications and validation were designed using Primers-4-Yeast (Yofe and Schuldiner, 2014). Primers and plasmids used are listed in Tables S2 and S3.

Yeast cells were grown overnight as a pre-culture in synthetic media (0.67% weight/volume yeast nitrogen base with ammonium sulphate, 2% weight/volume glucose, amino acid supplements, adenine-hemisulphate) at 30 °C, 280 rpm. Cells in glucose exhaustion (D exh.) were kept undiluted growing into stationary phase for 20-24 hours. For glucose replete conditions (High D), cells were back-diluted form the overnight culture and grown for 4 hours until reaching logarithmic growth in synthetic media with 2% glucose. Glucose restricted cells (D restr.) were collected from the overnight culture, washed twice with synthetic media containing 0.001% glucose and grown for 4 hours in this medium.

### Fluorescence microscopy

Cells were transferred to a 384 well glass bottom plate (Brooks) coated with Concanavalin A (Sigma-Aldrich). After 15 min of incubation, medium was replaced. Images were acquired either with an Olympus IX83 inverted fluorescence microscope with a Lumencor SpectraX LED light source and a 40x or 60x air objective using the Olympus ScanR Automated Image Acquisition Software or with an Opera Perkin Elmer microscope with a laser light source and a 60x water immersion objective (NA=1.2) using the Opera Software 2.0 (EvoShell). For Figure 1C-E cells were imaged using an iMIC-based microscope (FEI / Till Photonics) with an Olympus 100x oil objective (NA 1.45) and a laser light source. For staining of the vacuole, CellTracker Blue CMAC dye (7-amino-4-chloromethylcoumarin) (0.1 mM; ThermoFisher) was added to the cells for 30 minutes, followed by replacing of the medium and imaging.

### Library generation by automated mating and microscopy-based screening

The query strains Nvj1-Cherry and Zrc1-Cherry were constructed using a synthetic genetic array (SGA) compatible strain (Breslow et al., 2008) and crossed with a collection of strains selected from the genome wide SWAT mutant collection (Weill et al., 2018; Yofe et al., 2016) which consists of strains that are N-terminally GFP tagged using automated mating (Cohen and Schuldiner, 2011; Yan Tong and Boone, 2006; Yofe et al., 2016). Automated mating was performed using the RoToR benchtop colony array instrument (Singer instruments). Strains were mated on YPD medium (1% yeast extract (Thermo Fisher, 212750), 2% Bacto Peptone (Thermo Fisher, 211677), 2% Glucose (Sigma-Aldrich, G8270), 2.2% Agar (Becton Dickinson, 21030)) and selected for diploids before sporulation was induced by transfer to nitrogen starvation medium for 5 days. To select for haploids, cells were moved to plates containing 50 mg/L Canavanine (Sigma-Aldrich, C9758) and 50 mg/L Thialysine (Sigma-Aldrich, A2636). For the final selection step, cells were transferred to a plate containing selections for all desired mutations. For imaging the strains were grown in 384-well polystyrene plates in SD medium at the indicated growth condition. Cells were transferred to a 384-well glass plate (Brooks) using the Bench Smart 96^TM^ liquid handler (Mettler Toledo).

### Whole cell protein extraction, SDS-PAGE and western blotting

For the western blots 2.5 OD of cells in glucose replete (High D), glucose exhausted (D exh.) or glucose restriction (D restr.) condition were harvested by centrifugation and resuspended in Urea lysis buffer (8 M Urea, 50 mM Tris, pH 7.5) with protease cocktail (1:200, Merck). Acid washed glass beads were added, and the samples were vortexed at maximal velocity for 4x5 min, with 1 min pause on ice in between. The supernatant was collected, and a fraction was used to measure protein concentration by a Bradford assay. Then, cells were incubated for 5 min at 95°C in MSB ox. buffer (8 M Urea, 0.15% (w/v) bromphenol blue, 5 mM EDTA, 3.2% SDS, 100 mM Tris-HCl, 4% (v/v) glycerol, 10% β-mercaptoethanol). Samples of either cell extraction were subjected to SDS-PAGE (4–15% Mini-PROTEAN® TGX™ Precast Protein Gel, Bio-Rad) and subsequent western blotting. Antibodies used are listed in the Table S4. Signals were detected using the Lumi-Light ^PLUS^ substrate (Roche) and the Azure c600 imaging system. Quantification of protein levels in Figure 3G was performed with the AzureSpot software. Intensity of each band was measured and normalized to the respective GAPDH intensity. Ratio of Nvj1-Cherry compared to the control are depicted.

### High performance thin layer chromatography

Overnight cultures were grown in synthetic medium containing 2% glucose. For the logarithmic growth phase (High D) the same medium was used to inoculate the main cultures with an OD600 of 0.2. Cells were incubated at 30°C and shaken at 160 rpm until reaching an OD600 of 0.8. For glucose restriction an adequate amount of cells was collected from the overnight culture and washed twice with synthetic medium containing 0.001% glucose. The main culture, using the low glucose medium, was inoculated at an OD600 of 0.8. Cells were shaken at 160 rpm and 30°C for 4 h.

For lipid extraction a total of 180 mg of wet weight was harvested of each culture. The protocol for total lipid extraction was based on (Folch et al., 1957). 5 ml of a chloroform/methanol (CHCl_3_/MeOH) 2:1 (v/v) mixture and 1 ml of glass beads (Sigma-Aldrich, G8722) together with 125 µg of cholesteryl formate (Sigma Aldrich, S44853) as internal standard was added to each cell pellet. For lysis the mixture was shaken using a Heidolph Multi Reax shaker for 30 min at level 8. After addition of 1 ml H_2_O shaking was continued for another 10 min. The samples were centrifuged for 5 min at 2500 g which resulted in formation of two phases. The aqueous phase was carefully discarded, and 2 ml of an artificial upper phase comprised of MeOH/H_2_O/CHCl_3_ (48/47/3; v/v/v) was added. Following a short mixing, samples were again centrifuged (5 min, 2500 g), the aqueous phase discarded and the organic phase collected. A nitrogen stream was used to completely evaporate the solvent, and the dried samples were subsequently dissolved in 1 ml CHCl_3_/MeOH (2:1; v/v) and stored at -20°C.

For quantification of neutral lipid contents, between 30 and 100 µl of total lipid extracts were applied on a HPTLC silica gel 60 plate (20 x 10 cm) using a CAMAG automatic TLC sampler (CAMGA, ATS4). A mobile phase consisting of n-hexane/n-heptane/diethyl ether/acetic acid (63/18.5/18.5/1; v/v/v/v) was used to separate neutral lipids. This developing step was performed by the CAMAG automatic developing chamber (ADC2). To visualize the separated lipids the plate was derivatized with 0.01% primuline (Sigma-Aldrich, 206865) using the CAMAG derivatizer and afterwards heated to 40°C for 2 min on the CAMAG TLC plate heater 3. Image acquisition of the fully developed HPTLC plates was executed by the CAMAG TLC visualizer 2 using the VisionCATS software. To determine the different lipid concentrations, the HPTLC bands were converted into chromatograms and evaluated via a standard curve. This standard curve was generated by applying a neutral lipid standard mix together with the samples on the HPTLC plate in quantities ranging from 0.5 to 15 µg lipid. The standard mix was comprised of triacylglycerol ((15:0-18:1-15:0) Sigma-Aldrich, 330723C), diacylglycerol ((18:1) Sigma-Aldrich, 800811C), sodium oleate (Sigma-Aldrich, O7501), ergosterol (Thermo Fisher Scientific, 117810050), cholesteryl oleate ((18:1) Sigma-Aldrich,700269P) and cholesteryl formate (Sigma-Aldrich S448532) dissolved in CHCl_3_/MeOH (2:1; v/v) each at a concentration of 500 ng/µl. The neutral lipid contents were standardized according to the internal standard.

### Quantifications and statistical analysis

All microscopic images were processed using Fiji/ImageJ. Line graphs were generated using Fiji/ImageJ to generate an intensity profile after background subtraction. The intensity profiles were normalized to the maximum value and plotted using GraphPad Prism6.

For Pex31 localization in Figure 1F, cells were divided in three classes: (i) Pex31 signal that completely or partially colocalises with Nvj1-Cherry, (ii) Pex31 foci that are at the nuclear ER but not overlapping with Nvj1-Cherry signal and (iii) cells that do not show any Pex31 foci at the nuclear ER. Percentage of each class is depicted.

For quantification of enrichment of signal at the NVJ a first region of interest (ROI) was generated at the area defined as the NVJ. A second ROI was drawn either around the cell (Fig. 2C and D), opposite site of the nuclear ER (Fig. 2E and F) or on the cytosol (Fig. 2G). The ratio between the mean signal at the NVJ versus the control ROI is depicted. For measurement of the NVJ area (Fig. 3B, 3D), ROIs of the cells to be analyzed were generated and overlayed on a binary image in which the NVJ was selected using the Otsu threshold. The size of the NVJ was then measured using the analyze particle function of ImageJ. For vacuolar fragmentation (Fig. 4B), vacuoles in each cell were counted and cells were categorized into either one vacuole (1 vac.) or two or more vacuoles (≥2 vac.) containing cells. For vacuolar domains, z-stacks of Vph1-mKate2 expressing cells were imaged (Fig. 4C). Based on the z-stacks each cell was categorized dependent on vacuolar domain content.

For visualization and statistic comparison Graph Pad Prism 6 was used. The bar graphs in Figure 3H and 4D were generated using Graph Pad Prism 10.

Data was obtained from at least three independent experiments. The number of cells analyzed is indicated in the respective figure legend. The whiskers of box plots are indicated in the figure legends. For area measurement the whiskers depict 5-95% of the data. For comparison between two strains a t-test was used (Fig. 2C, 2D, 2E, 2F, 2G, 4B, 4C). Comparison of multiple samples was performed using one-way ANOVA with Tukeys multiple comparisons (Fig. 3B, 3D, 3F, 3J, 3L) or Bonferroni (Figure 4D). Statistical significance was defined as * p<0.05; **, p<0.01; ***, p<0.001.

## ACKNOWLEDGEMENTS

M.H. and D.T.V.D. are members of CiM-IMPRS, the joint graduate school of the Cells-in-Motion Interfaculty Centre, University of Münster, Germany and the International Max Planck Research School – Molecular Biomedicine, Münster, Germany. We thank Maya Schuldiner and Roland Wedlich-Söldner for discussions, strains and plasmids, Christian Schuberth for insightful comments on image analysis and quantifications, and all members of the Bohnert lab for discussions.

## COMPETING INTERESTS

No competing interests declared.

## FUNDING

This work was supported by the Gerty Cori Programme, Medical Faculty, University of Münster (to M.B.), and the Deutsche Forschungsgemeinschaft (DFG, German Research Foundation), SFB1190 P21 (project ID 264061860), SFB1348 A13 (project ID 386797833), SFB1557 P03 (project ID 467522186), and FOR5815 P6 (project ID 538651361) (to M.B.).

## AUTHOR CONTRIBUTION

Conceptualization: M.H., M.B.; Investigation and Formal Analysis: M.H., P.H., X.-T.Z., M.W., D.T.V.D., R.M.F.; Visualization: M.H., P.H., R.M.F., MB; Writing – Original Draft: M.H., M.B.; Writing – Review and editing M.H., P.H., X.-T.Z., M.W., D.T.V.D., R.M.F., MB; Supervision and Funding Acquisition: M.B.

## DATA AND RESOURCE AVAILABILITY

Requests for resources and reagents should be directed to Maria Bohnert.

## Notes

### Competing Interest Statement

The authors have declared no competing interest.

