## Supplement for "Role of Pex31 in metabolic adaptation of the nucleus vacuole junction NVJ"

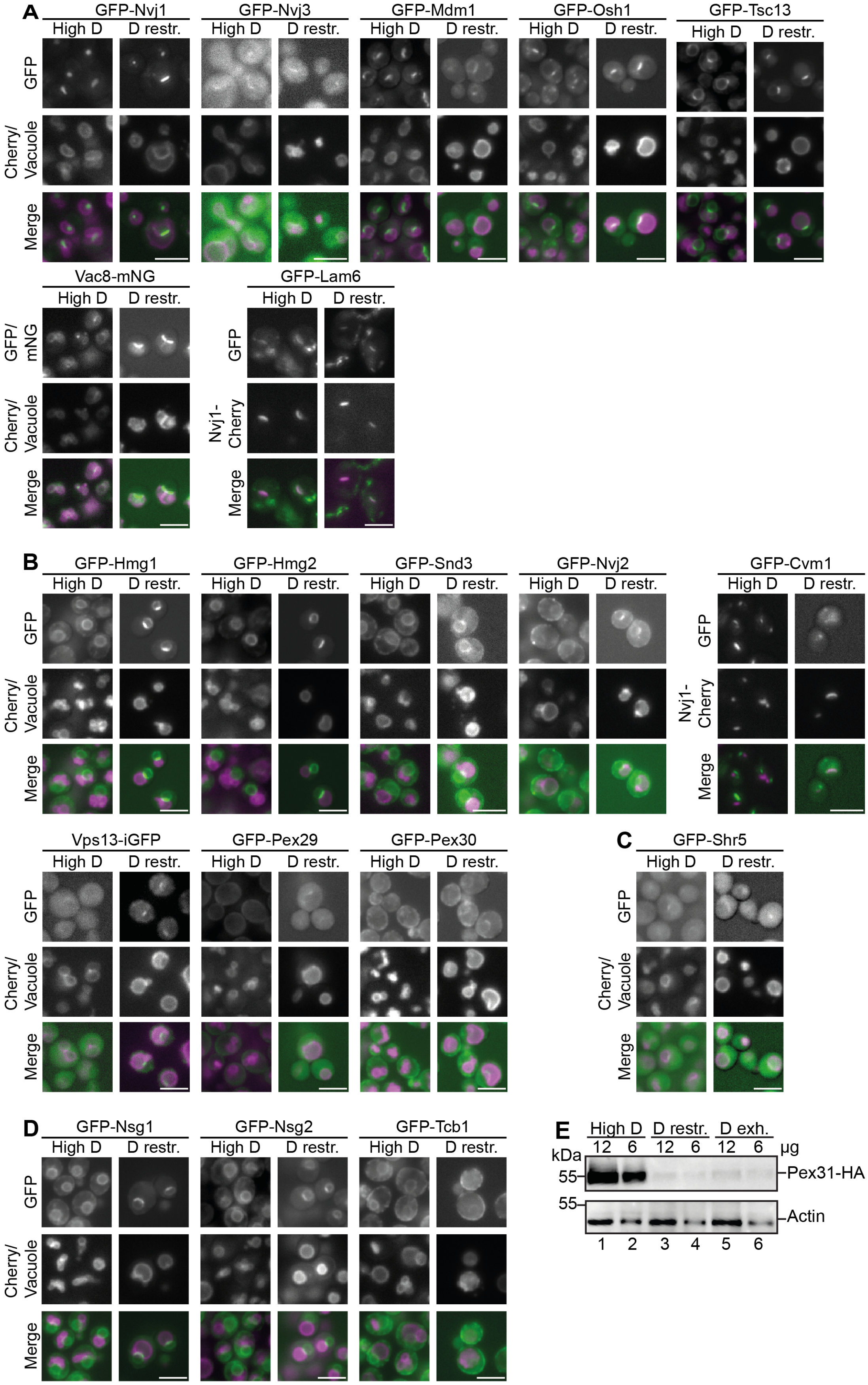
A
GFP-Nvj1
GFP-Nvj3
GFP-Mdm1
GFP-Osh1
GFP-Tsc13
High D
D restr.
GFP
Cherry/ Vacuole
Merge
Vac8-mNG
GFP-Lam6
GFP/ mNG
Nvj1-Cherry
B
GFP-Hmg1
GFP-Hmg2
GFP-Snd3
GFP-Nvj2
GFP-Cvm1
Vps13-iGFP
GFP-Pex29
GFP-Pex30
C
GFP-Shr5
D
GFP-Nsg1
GFP-Nsg2
GFP-Tcb1
E
High D
D restr.
D exh.
12
6
12
6
12
6
µg
kDa
55
55
Pex31-HA
Actin
1
2
3
4
5
6

**Figure S1**

**Figure S1.** Related to Figure 1. A microscopy-based screen identifies permanent and conditional residents of the nucleus vacuole junction NVJ.

(A)-(D) Original images of hits identified in screen described in Fig. 1A and B. Using automated mating, either Nvj1-Cherry or Zrc1-Cherry were introduced into a collection of strains expressing proteins related to lipid handling fused with green fluorescent tags. Strains were cultured overnight on medium containing 2% glucose and either back diluted in fresh medium containing 2% glucose and grown to logarithmic growth phase (High D), or cells were back diluted in medium containing 0.001% glucose and incubated for four hours (D restr.). All strains were analyzed by automated microscopy and classified as permanent (A and C) or conditional (B and D) NVJ residents. Scale bars, 5 µm.

(E) Pex31-HA cells were cultured at indicated conditions and subjected to SDS-PAGE and western blotting.

**Table S1.** List of yeast strains used in this study.

| Name | Genotype | Source | Identifier |
|---|---|---|---|
| WT | his3Δ1 leu2Δ0 lys2+/lys+ met15Δ0 ura3Δ0 can1Δ::STE2pr-sp HIS5 lyp1Δ::STE3pr-LEU2 | Breslow et al., 2008 | yMB3 |
| Nvj1-Cherry GFP-Shr5 | his3Δ1 leu2Δ0 lys2+/lys+ met15Δ0 ura3Δ0 can1Δ::STE2pr-sp HIS5 lyp1Δ::STE3pr-LEU2 Nvj1-Cherry::NAT pNOP1-GFP-Shr5 | This study | yMB1276 |
| Nvj1-Cherry GFP-Nsg1 | his3Δ1 leu2Δ0 lys2+/lys+ met15Δ0 ura3Δ0 can1Δ::STE2pr-sp HIS5 lyp1Δ::STE3pr-LEU2 Nvj1-Cherry::NAT pNOP1-GFP-Nsg1 | This study | yMB1275 |
| Nvj1-Cherry GFP-Nsg2 | his3Δ1 leu2Δ0 lys2+/lys+ met15Δ0 ura3Δ0 can1Δ::STE2pr-sp HIS5 lyp1Δ::STE3pr-LEU2 Nvj1-Cherry::NAT pNOP1-GFP-Nsg2 | This study | yMB1273 |
| Nvj1-Cherry GFP-Tcb1 | his3Δ1 leu2Δ0 lys2+/lys+ met15Δ0 ura3Δ0 can1Δ::STE2pr-sp HIS5 lyp1Δ::STE3pr-LEU2 Nvj1-Cherry::NAT pNOP1-GFP-Tcb1 | This study | yMB1274 |
| Nvj1-Cherry Pex31-mNG | his3Δ1 leu2Δ0 lys2+/lys+ met15Δ0 ura3Δ0 can1Δ::STE2pr-sp HIS5 lyp1Δ::STE3pr-LEU2 Nvj1-mCherry::NAT Pex31-mNG::HYG | This study | yMB1090 |
| Δ*pex31* | his3Δ1 leu2Δ0 lys2+/lys+ met15Δ0 ura3Δ0 can1Δ::STE2pr-sp HIS5 lyp1Δ::STE3pr-LEU2 Δpex31::G418 | This study | yMB273 |
| GFP-NVJ2 | his3Δ1 leu2Δ0 lys2+/lys+ met15Δ0 ura3Δ0 can1Δ::STE2pr-sp HIS5 lyp1Δ::STE3pr-LEU2 pNop1-GFP-Nvj2::URA | This study | yMB1001 |
| GFP-NVJ2 Δ*pex31* | his3Δ1 leu2Δ0 lys2+/lys+ met15Δ0 ura3Δ0 can1Δ::STE2pr-sp HIS5 lyp1Δ::STE3pr-LEU2 Δpex31::G418 pNop1-GFP-Nvj2::URA | This study | yMB1002 |
| Nvj1-Cherry Nsg1-GFP | his3Δ1 leu2Δ0 lys2+/lys+ met15Δ0 ura3Δ0 can1Δ::STE2pr-sp HIS5 lyp1Δ::STE3pr-LEU2 Nvj1-mCherry::Nat Nsg1-GFP::His | This study | yMB1011 |
| Nvj1-Cherry Δ*pex31* Nsg1-GFP | his3Δ1 leu2Δ0 lys2+/lys+ met15Δ0 ura3Δ0 can1Δ::STE2pr-sp HIS5 lyp1Δ::STE3pr-LEU2 ΔPex31::G418 Nvj1-mCherry::Nat Nsg1-GFP::His | This study | yMB1013 |
| Nvj1-Cherry Hmg2-GFP | his3Δ1 leu2Δ0 lys2+/lys+ met15Δ0 ura3Δ0 can1Δ::STE2pr-sp HIS5 lyp1Δ::STE3pr-LEU2 Nvj1-mCherry::Nat Hmg2-GFP::His | This study | yMB1010 |
| Nvj1-Cherry Hmg2-GFP Δpex31 | his3Δ1 leu2Δ0 lys2+/lys+ met15Δ0 ura3Δ0 can1Δ::STE2pr-sp HIS5 lyp1Δ::STE3pr-LEU2 Δpex31::G418 Nvj1-Cherry::NAT Hmg2-GFP::HIS3MX6 | This study | yMB1020 |
| Pex29-mNG Nvj1-Cherry | his3Δ1 leu2Δ0 lys2+/lys+ met15Δ0 ura3Δ0 can1Δ::STE2pr-sp HIS5 lyp1Δ::STE3pr-LEU2 Pex29-mNG::HYG Nvj1-mCherry::NAT | This study | yMB1117 |
| Pex29-mNG Nvj1- | his3Δ1 leu2Δ0 lys2+/lys+ met15Δ0 ura3Δ0 can1Δ::STE2pr-sp HIS5 lyp1Δ::STE3pr- | This study | yMB1118 |

| | | | |
|---|---|---|---|
| mCherry Δ*pex31* | LEU2 Pex29-mNG::HYG Nvj1-mCherry::NAT Δpex31::G418 | | |
| Nvj1-Cherry Pex30-mNG Δpex31 | his3Δ1 leu2Δ0 lys2+/lys+ met15Δ0 ura3Δ0 can1Δ::STE2pr-sp HIS5 lyp1Δ::STE3pr-LEU2 Pex30-mNG::HYG Nvj1-mCherry::NAT Δpex31::G418 | This study | yMB1188 |
| Nvj1-Cherry Pex30-mNG | his3Δ1 leu2Δ0 lys2+/lys+ met15Δ0 ura3Δ0 can1Δ::STE2pr-sp HIS5 lyp1Δ::STE3pr-LEU2 Pex30-mNG::HYG Nvj1-mCherry::NAT | This study | yMB1159 |
| Nvj1-Cherry | his3Δ1 leu2Δ0 lys2+/lys+ met15Δ0 ura3Δ0 can1Δ::STE2pr-sp HIS5 lyp1Δ::STE3pr-LEU2 Nvj1-Cherry::NAT | This study | yMB908 |
| Nvj1-Cherry Δ*pex31* | his3Δ1 leu2Δ0 lys2+/lys+ met15Δ0 ura3Δ0 can1Δ::STE2pr-sp HIS5 lyp1Δ::STE3pr-LEU2 Δpex31::G418 Nvj1-Cherry::NAT | This study | yMB900 |
| Nvj1-Cherry Δ*snd3* | his3Δ1 leu2Δ0 lys2+/lys+ met15Δ0 ura3Δ0 can1Δ::STE2pr-sp HIS5 lyp1Δ::STE3pr-LEU2 Δsnd3::HYG Nvj1-mCherry::NAT | This study | yMB998 |
| Nvj1-cherry Δ*snd3* Δ*pex31* | his3Δ1 leu2Δ0 lys2+/lys+ met15Δ0 ura3Δ0 can1Δ::STE2pr-sp HIS5 lyp1Δ::STE3pr-LEU2 Nvj1-mCherry::G418 Δsnd3::NAT Δpex31::G418 | This study | yMB1115 |
| Snd3-GFP | his3Δ1 leu2Δ0 lys2+/lys+ met15Δ0 ura3Δ0 can1Δ::STE2pr-sp HIS5 lyp1Δ::STE3pr-LEU2Snd3-GFP::NAT | This study | yMB1119 |
| Snd3-GFP Δ*pex31* | his3Δ1 leu2Δ0 lys2+/lys+ met15Δ0 ura3Δ0 can1Δ::STE2pr-sp HIS5 lyp1Δ::STE3pr-LEU2 Snd3-GFP::NAT Δpex31::G418 | This study | yMB1120 |
| Pex30-mNG Δ*snd3* | his3Δ1 leu2Δ0 lys2+/lys+ met15Δ0 ura3Δ0 can1Δ::STE2pr-sp HIS5 lyp1Δ::STE3pr-LEU2 Pex30-mNG::NAT Δsnd3::HYG | This study | yMB1133 |
| Pex30-mNGn Δ*snd3*Δpex31 | his3Δ1 leu2Δ0 lys2+/lys+ met15Δ0 ura3Δ0 can1Δ::STE2pr-sp HIS5 lyp1Δ::STE3pr-LEU2 Pex30-mNG::NAT Δpex31::G418 Δsnd3::HYG | This study | yMB1134 |
| Pex31-HA | his3Δ1 leu2Δ0 lys2+/lys+ met15Δ0 ura3Δ0 can1Δ::STE2pr-sp HIS5 lyp1Δ::STE3pr-LEU2 Pex31-HA::HIS | This study | yMB302 |
| Zrc1-Cherry | his3Δ1 leu2Δ0 lys2+/lys+ met15Δ0 ura3Δ0 can1Δ::STE2pr-sp HIS5 lyp1Δ::STE3pr-LEU2Snd3-GFP::NAT Zrc1-Cherry::NAT | This study | yMB368 |
| Vph1-mkate2 | his3Δ1 leu2Δ0 lys2+/lys+ met15Δ0 ura3Δ0 can1Δ::STE2pr-sp HIS5 lyp1Δ::STE3pr-LEU2 Vph1-mKate2::NAT | This study | yMB813 |
| Vph1-mKate2 Δ*pex31* | his3Δ1 leu2Δ0 lys2+/lys+ met15Δ0 ura3Δ0 can1Δ::STE2pr-sp HIS5 lyp1Δ::STE3pr-LEU2 Vph1-mKate2::NAT Δpex31::G418 | This study | yMB1114 |

**Table S2.** List of plasmids used in this study.

| Name | Utilization | Source | Identifier |
|---|---|---|---|
| pFA6a-KanMX6 | Yeast genomic manipulation (deletion) | Longtine et al., 1998 | pMB5 |

| | | | |
|---|---|---|---|
| pFA6a-NatMX6 | Yeast genomic manipulation (deletion) | Goldstein and McCusker, 1999 | pMB6 |
| pFA6-Hygro | Yeast genomic manipulation (deletion) | Goldstein and McCusker, 1999 | pMB10 |
| pFA6a-HIS3 | Yeast genomic manipulation (deletion) | Longtine et al., 1998 | pMB49 |
| pFA6a-NAT-Cherry | Yeast genomic manipulation (N-terminal tagging with Cherry) | Longtine et al., 1998 | pMB9 |
| pFA6a-NAT-eGFP | Yeast genomic manipulation (N-terminal tagging with eGFP) | Longtine et al., 1998 | pMB12 |
| pYM2 3HA-HIS3MX6 | Yeast genomic manipulation (C-terminal tagging with 3xHA) | Janke et al., 2004 | pMB230 |
| pYM25-5xGA-mNeonGreen-HygR | Yeast genomic manipulation (C-terminal tagging with mNG) | Haase et al., 2023 | pMB451 |
| pYM42-5xGA-mNeonGreen-NatR | Yeast genomic manipulation (C-terminal tagging with mNG) | Haase et al., 2023 | pMB91 |

**Table S3.** List of primers used in this study.

| Primer name | Sequence | Identifier |
|---|---|---|
| pMS80 chk F | ggcatggacgagctgtacaag | prMB22 |
| pFA6 F1 rev com | TTAATTAACCCGGGGATCCG | prMB23 |
| PEX31 5'UTR CHK F | TAATGTGGAGTGGGTAATCG | prMB162 |
| PEX31 KO pFA6 F | CTGGTTGTCAAGCCTTGGTTTCCCTTTATTTGATAG TATGcggatccccgggttaattaa | prMB163 |
| PEX31 KO pFA6 R | AGTGTGAACGTTGTTGTCCATATGGGGCATGCACT CATTAgaattcgagctcgtttaaac | prMB164 |
| PEX31 WT CHK F | GAATGGCACGATAAAGATTG | prMB165 |
| PEX31 WT CHK R | TACTGTCCGAGGAAGGTATG | prMB166 |
| Nvj1-Ctag-chk_F | TAAGGACATGAACGTTTTGG | prMB829 |
| Nvj1-Ctag-pFA6_F | AGTGAACACTGAACAAGCATACTCTCAACCATTTAG ATACcggatccccgggttaattaa | prMB830 |
| Nvj1-Ctag-pFA6_R | GTGACGATGATAACCGAGATGACGGAAATATAGTA CATTAgaattcgagctcgtttaaac | prMB831 |
| PEX31 C-tag CHK F | GAATGGCACGATAAAGATTG | prMB186 |
| PEX31 C-tag pYM F | ATTAATACAAATATCTGATGTTTCAATGTCTCCTTCT CTAcgtacgctgcaggtcgac | prMB187 |

| | | |
|---|---|---|
| PEX31 C-tag pYM R | AGTGTGAACGTTGTTGTCCATATGGGGCATGCACT CATTAatcgatgaattcgagctcg | prMB188 |
| NOP1-NVJ2-gDNA-F | CTTTCATTTTTGCACCTAATTGGTATGGCACATTTTT CAGTTTTCTCCCACTTAAGTTTTC | prMB1491 |
| NOP1-NVJ2-gDNA-R | TCTGCTGTATCATGATCTAGCTCCTTCTTCGATGCC GATTTAAGGTTACTGTACAATAAA | prMB1492 |
| NVJ2 N-tag CHK R | GTTCTTTGCAAGATTTGCTC | prMB1493 |
| HMG2_Ctag_CHK_F | ATTTGGTCACTGCACTTTTC | prMB1496 |
| HMG2_Ctag_pFA6_F | AAGTAACAAAGGGCCCCCCTGTAAAACCTCAGCAT TATTAcggatccccgggttaattaa | prMB1497 |
| HMG2_Ctag_pFA6_R | ACAAAGATATAAAGTATCACCATGTAAACTACAAGA GTTAgaattcgagctcgtttaaac | prMB1498 |
| NSG1_Ctag_CHK_F | ACCATTCATTGACTCTTTCG | prMB1499 |
| NSG1_Ctag_pFA6_F | GTTTTTGATGTTCAGCAAGTTGGGCAGATATTTATT CAATcggatccccgggttaattaa | prMB1500 |
| NSG1_Ctag_pFA6_R | CATCGATACTAATCATTGAACGCCCCTATGGGAACA CTTAgaattcgagctcgtttaaac | prMB1501 |
| PEX29 C-tag CHK F | GGATCCAAAAGAATGGGTAG | prMB1775 |
| PEX29 C-tag pYM F | GTCAATCGAAGAGCTAACAGACACTCTCAATTCAAC TATAcgtacgctgcaggtcgac | prMB1776 |
| PEX29 C-tag pYM R | TGTATCATCAGTGAACATATAGTATAACAAATCAAG TTTAatcgatgaattcgagctcg | prMB1777 |
| PEX30 C-tag CHK F | GGTAAGAACCGCAGAATTG | prMB182 |
| PEX30 C-tag pYM F | ATCAAATCCAACCATTGGTCGCGATAGCAAGAAGG CCGTAcgtacgctgcaggtcgac | prMB183 |
| PEX30 C-tag pYM R | AGATTATATTATGTAAAGGTAAAAACGGGAGCGAG CATCAatcgatgaattcgagctcg | prMB184 |
| SND3 C-tag CHK F | TCTTCCATCGGTATCTCTTG | prMB373 |
| SND3 C-tag pFA6 F | AGAAGCTGAAAGAGCCGGTAACGCTGGTGTTAAGG CTGAAcggatccccgggttaattaa | prMB374 |
| SND3 C-tag pFA6 R | AAAACTAGGAAAAAAAAATACTTCGCTTTTGATCGA ATCAgaattcgagctcgtttaaac | prMB375 |
| VPH1-C-tag-pFA6_F | GGAAGTCGCTGTTGCTAGTGCAAGCTCTTCCGCTT CAAGCcggatccccgggttaattaa | prMB1002 |
| VPH1-C-tag-pFA6_R | ACTTAAATGTTTCGCTTTTTTTAAAAGTCCTCAAAAT TTAgaattcgagctcgtttaaac | prMB1003 |

**Table S4.** List of antibodies used in this study.

| **Antibodies** | **Source** | **Identifier** |
|---|---|---|
| Cherry | ThermoFisher | PA5-34974 |
| GAPDH | Abcam | Cat#125247; RRID: AB_11129118 |
| HA | Proteintech | Cat# 51064-2-AP |
| Actin | MP Biomedicals | 08691001 |
| anti-mouse IgG | Dianova | Cat#115-035-003 |
| anti-rabbit IgG | Dianova | Cat#111-035-003 |
